# Single cell expression analysis uncouples transdifferentiation and reprogramming

**DOI:** 10.1101/351957

**Authors:** Mirko Francesconi, Bruno Di Stefano, Clara Berenguer, Marisa de Andres, Maria Mendez Lago, Amy Guillaumet-Adkins, Gustavo Rodriguez-Esteban, Marta Gut, Ivo G. Gut, Holger Heyn, Ben Lehner, Thomas Graf

## Abstract

Many somatic cell types are plastic, having the capacity to convert into other specialized cells (transdifferentiation)(*1*) or into induced pluripotent stem cells (iPSCs, reprogramming)(*2*) in response to transcription factor over-expression. To explore what makes a cell plastic and whether these different cell conversion processes are coupled, we exposed bone marrow derived pre-B cells to two different transcription factor overexpression protocols that efficiently convert them either into macrophages or iPSCs and monitored the two processes over time using single cell gene expression analysis. We found that even in these highly efficient cell fate conversion systems, cells differ in both their speed and path of transdifferentiation and reprogramming. This heterogeneity originates in two starting pre-B cell subpopulations,large pre-BII and the small pre-BII cells they normally differentiate into. The large cells transdifferentiate slowly but exhibit a high efficiency of iPSC reprogramming. In contrast, the small cells transdifferentiate rapidly but are highly resistant to reprogramming. Moreover, the large B cells induce a stronger transient granulocyte/macrophage progenitor (GMP)-like state, while the small B cells undergo a more direct conversion to the macrophage fate. The large cells are cycling and exhibit high Myc activity whereas the small cells are Myc low and mostly quiescent. The observed heterogeneity of the two cell conversion processes can therefore be traced to two closely related cell types in the starting population that exhibit different types of plasticity. These data show that a somatic cell’s propensity for either transdifferentiation and reprogramming can be uncoupled.

**One sentence summary:** Single cell transcriptomics of cell conversions

## Main Text

C/EBPα is a master regulator of myelopoiesis(*3*). When overexpressed in B cell precursors, it induces their efficient transdifferentiation into macrophages(*1*), and when transiently overexpressed, it poises them for rapid and highly efficient reprogramming into iPSCs in response to induction of Oct4, Sox2, Klf4 and Myc (OSKM)(*4*). The combination of these two systems gives us the unique opportunity to study the determinants of both types of cell conversion by following gene expression in single cells starting from the same cell population.

We isolated CD19^+^ B cells precursors from the bone marrow of reprogrammable mice(*5*) and infected them with a retrovirus encoding a hormone inducible form of C/EBPa (Cebpa-ER-hCD4). After expansion in culture, we induced them to either trans-differentiate into macrophages or to reprogram into iPSCs. To induce the macrophage fate, we treated the cells with beta-estradiol (E2) to activate C/EBPα. To induce the iPSC fate, we treated the cells with E2 for 18 hours, washed out the hormone and added doxycycline to induce OSKM(*4, 6*). For transdifferentiation, we collected cells before (0h) and after 6h, 18h, 42h, 66h and 114h of C/EBPα induction; for reprogramming samples were prepared at days 2, 4, 6 and 8 after OSKM induction of 18h C/EBPa-pulsed cells (Fig. 1A). We collected two pools of 192 cells at each time point and sequenced their RNA using MARS-Seq(*7*). After quality control and filtering (see Methods), we obtained expression profiles for 17,183 genes in 3,152 cells. We then performed dimensionality reduction, corrected for batch effects, and extracted gene expression signatures (See Methods and Fig. S1-3, Table S1-4). Visualizing the data using diffusion maps(*8*) revealed branching between transdifferentiation and reprogramming at the 18h time-point, with largely synchronous cohorts of cells moving along distinct trajectories and reaching homogenous final cell populations consisting of either induced macrophage (iMac) or iPSC-like cells, respectively (Fig. 1B). We observed no branching into alternative routes, in contrast to what has been described for the transdifferentiation of fibroblasts into neurons(*9*), muscle cells(*10*) or iPSCs(*11, 12*).

**Fig. 1.**
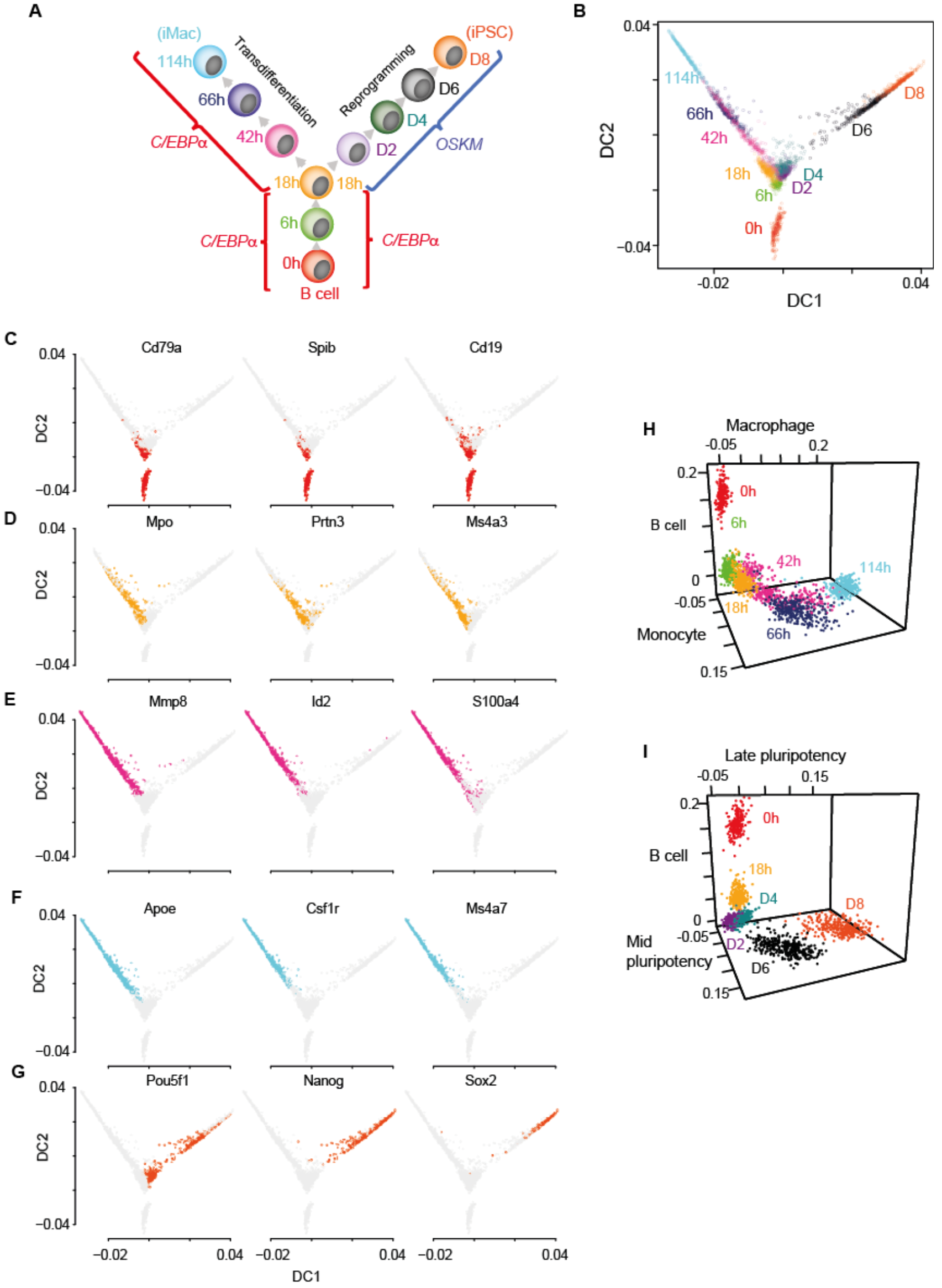
Single cell gene expression analysis of B cell to macrophage transdifferentiation and B cell to iPSC reprogramming. **A,** Overview of the experimental design, showing time points analysed. **B,**Single cell projections onto the first two diffusion components (DC1 and DC2). **C-F**, as in **B,** with top 50% of cells expressing selected markers for B cells in red (**C**), GMP/granulocytes in orange (**D**), monocytes in purple (**E**), macrophages in light blue (**F**) and pluripotent cells in orange-red (**G**). **H-I,** Projection of transdifferentiating cells onto B cell, macrophage, and monocyte specific independent components (**H**), and reprogramming cells onto, B cell, mid-and late - pluripotency specific independent components (**I**).

B cell genes become largely silenced after 18h (Fig. 1C and Fig. S4A). During transdifferentiation, there is a transient activation of granulocyte/GMP genes (Fig. 1D, Fig. S4B), followed by activation of monocyte (Fig. 1E, Fig. S4C) and then macrophage genes (Fig. 1F, Fig. S4D). After OSKM induction, endogenous *Pou5f1* (*Oct4*) is activated at day 2, followed by expression of *Nanog* at day 4 and *Sox2* at day6 (Fig. 1G, Fig. E-H), consistent with the high reprogramming efficiency of our system(*6, 13*).

Visualizing single cells in the expression space spanned by B cell, monocyte and macrophage programs (Fig. 1H) and B cell, mid and late reprogramming (Fig. 1I), however, reveals a degree of asynchrony. To identify potential causes of this asynchronous behaviour, we determined which independent component analysis (ICA)-derived expression signatures (Fig. S2) best predicted cell progression toward the macrophage state (Fig. 2A) at each time-point (excluding expression signatures directly involved in transdifferentiation, that is the B cell, monocyte, granulocyte, and macrophage programs). We found that a signature highly enriched in Myc target genes (component 5, Fig. S2, Table S4) best predicts and negatively correlates with the extent of transdifferentiation at intermediate time points (Fig. 2B, Fig. S5). Expression of the Myc targets varies extensively across cells within each time point but changes little during transdifferentiation (Fig. 2C). The data therefore suggest that the cells with lower expression of Myc targets transdifferentiate more rapidly into macrophages.

**Fig. 2.**
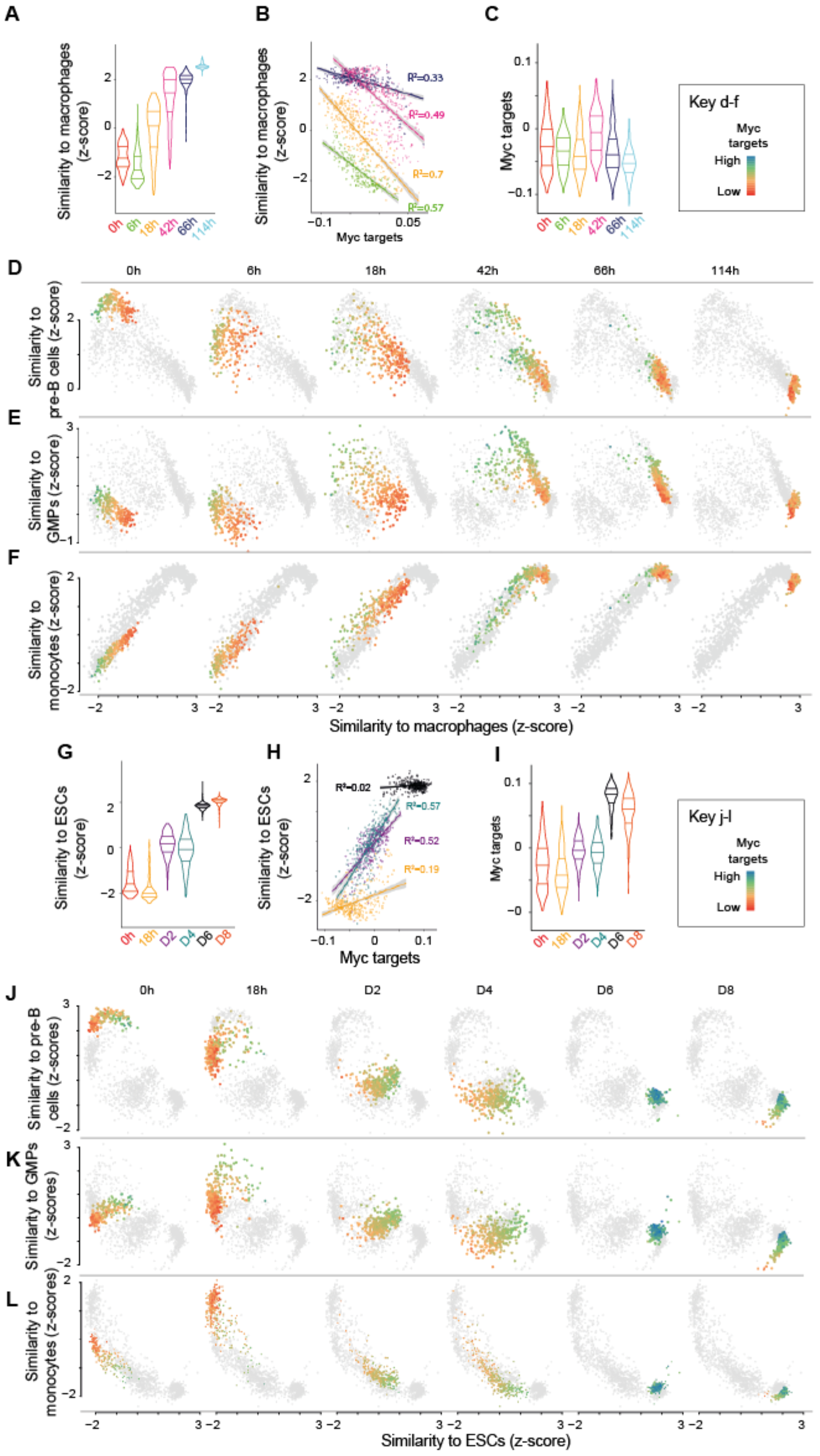
Myc target levels predict differences in single cell transdifferentiation and reprogramming trajectories. **A,** Distribution of gene expression similarity between single cells and reference bone marrow derived macrophages(*26*) (acquisition of macrophage state) during transdifferentiation. **B,**Correlation between the Myc targets component and acquisition of macrophage state from **A**. **C**, Expression of Myc targets at the various transdifferentiation time points. **D-F,**Single cell trajectories relating the B cell state (**D**), the GMP state (**E**) and the monocyte state (**F**) to the acquisition of the macrophage state during transdifferentiation. **G,**Distribution of expression similarity between single cells and reference embryonic stem cells (ESCs) during reprogramming. **H**, Correlation between Myc targets and acquisition of pluripotency from **G**. **I,**Expression of Myc targets at the various reprogramming time points. **J**-**L**as in **D-F,**but relating differentiation states with acquisition of the pluripotent state (ESCs) (see also Figure S8).

We next tested how the expression of Myc targets relates to the loss of the B cell state and the acquisition of transient myeloid-like cell states during transdifferentiation.

Visualizing similarity to the pre-B cell state shows that low expression of Myc targets is more strongly associated with a rapid gain of the macrophage state than with a rapid loss of the B cell state (Fig. 2D). Moreover, higher Myc target expression is associated with a larger and more persistent induction of a GMP-like state (Fig. 2E). Myc target expression does not associate with the extent of induction of a transient monocyte-(Fig. 2F) or granulocyte-like state (Fig. S6). In conclusion, cells with low expression of Myc targets acquire the macrophage fate more rapidly and transdifferentiate via a less pronounced transient induction of a GMP-like state.

We similarly searched for expression signatures that predict the progression of individual cells toward pluripotency within each time-point during reprogramming to iPSCs (Figure 2g). The expression of Myc targets was again predictive of cell fate conversion especially at early stages, however, in contrast to what was observed during transdifferentiation, high expression of Myc targets is associated with a more advanced state of reprogramming (Fig. 2H, Fig. S7). Moreover – and also different to what was observed during transdifferentiation – the expression of Myc targets increases during reprogramming (Fig. 2I). Visualizing similarity to pre-B cells, GMPs and monocytes during reprogramming shows that cells with high expression of Myc targets and a transient GMP state are at the forefront of the reprogramming trajectory. In contrast, cells with low expression of Myc targets lag behind and retain the monocyte program at D4 (Fig. 2J-L).

Is the heterogeneity in the expression of Myc targets due to a differential response of pre-B cells to the lineage instructive transcription factors or does it reflect a heterogeneity in the starting cell population? Examining the uninduced pre-B cells reveals substantial variation in the expression of Myc targets (Fig. 3A), suggesting that the heterogeneity pre-exists in the starting cell population. This variation also correlates with higher expression of both G1/S and G2/M phase cell cycle genes (Fig. 3A).

**Fig. 3.**
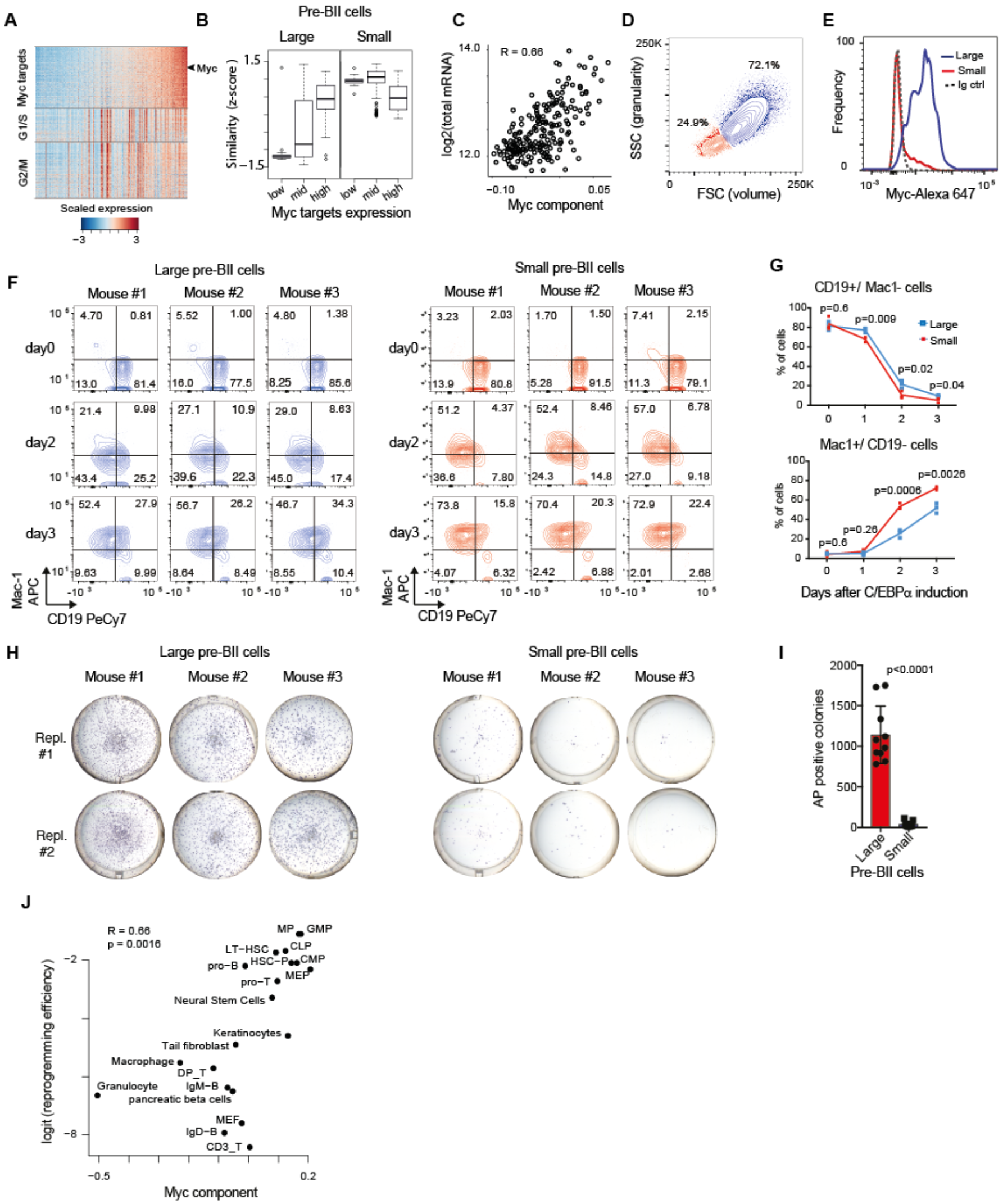
Two types of pre-B cells exhibit distinct cell conversion plasticities. **A**, Heatmap showing the expression of Myc targets, G1/S and G2/M specific genes in the starting pre-B cells sorted by Myc targets component. **B**, Similarity score of single cells binned by Myc targets expression (bottom 20%, mid and top 20%) with reference large and small pre-BII cells. **C**, Correlation between total mRNA molecules per cell and Myc targets expression. **D**, Representative FACS plot of starting pre-B cells showing forward (FSC) and side scatter (SSC). **E**, Representative FACS analysis of Myc levels detected in the 30% largest and the 30% smallest pre-B cell fractions. **F**, FACS plots of myeloid marker (Mac-1) and B cell marker (CD19) expression during induced transdifferentiation of sorted large and small pre-BII cells. **G**, Quantification of the results shown in **F** (n=3, Statistical significance was determined using multiple t-test with 1% false discovery rate). **H,**Visualization of iPSC-like colonies (stained by alkaline phosphatase) 12 days after OSKM induction of sorted large and small pre-BII cells. **I**, Quantification of the results shown in **H** (Statistical significance was determined using a two-tailed unpaired Student’s t-test). **J** Scatterplot showing the correlation between Myc component expression in different starting cell types (x-axis) and their corresponding (logit transformed) reprogramming efficiency (y-axis).

During B cell development in the bone marrow, large pre-BII cells undergo a proliferation burst and differentiate into quiescent small pre-BII cells(*14*), via Bcl6 induced transcriptional repression of *Myc*, events that are required for the initiation of light chain immunoglobulin rearrangements(*15*). Thus, heterogeneity in the starting pre-B cell population likely reflects variability along this B cell developmental transition. Comparing our single cell data with bulk expression data of cells at various stages of B cell development(*14, 16*) supports this hypothesis, showing that cells with higher expression of Myc targets are more similar to large pre-BII cells or cycling pre-B cells, while cells with lower expression of Myc targets are more similar to small pre-BII cells and non-cycling pre-B cells (Fig. 3B, Fig. S9A). Indeed, total mRNA content in our single pre-B cells varies within a three-fold range and scales with the expression of Myc targets, further suggesting a Myc dependent heterogeneity in cell size in the starting cell population (Fig. 3C).

Taken together, these observations suggest a pre-existing variation in the starting cell population, reflecting the developmental transition from large to small pre-BII cells and that this heterogeneity affects the speed of transdifferentiation and reprogramming in reciprocal ways: small pre-BII cells transdifferentiate faster but reprogram slowly, while large pre-BII cells transdifferentiate slowly but reprogram faster.

To further test this model, we analyzed our starting pre-B cell population by flow cytometry and found that it can be resolved by size and granularity into two discrete subpopulations, with about 1/3 small and 2/3 large cells (Fig. 3D). Intracellular staining of Myc monitored by flow cytometry confirmed that the larger cells express Myc while the smaller cells are Myc negative (Fig. 3E, Figs. S9B, S10A). These two subpopulations show the predicted differences for large and small pre-BII cells in cell proliferation(*15*), with the large cells incorporating 400 times more EdU within 2 hours than the small cells, showing that they proliferate while the small cells are largely quiescent (Figs. S9C, S10B). To determine whether the two cell types differ in their plasticity, we isolated B cell progenitors from reprogrammable mice and tested their ability to transdifferentiate and reprogram. In response to a continuous exposure to C/EBPa, the small pre-BII cells upregulated the macrophage marker Mac-1 faster and downregulated CD19 slightly more rapidly than large pre-BII cells (Figs. 3F-G, Fig. S10C). Similarly, the small cells acquired higher granularity, a marker of mature myeloid cells, and also slightly increased in volume compared to the large cells (Figs. S9D, S10D). In stark contrast, when 18h pulsed cells (also designated ‘Ba cells’(*4, 6*) were tested for reprogramming in response to OSKM induction, large pre-BII cells generated 30x times more iPSC colonies than small pre-B cells (Figs. 3H-I). Previous work has shown that C/EBPa induced maximal reprogramming efficiency by 18hrs of treatment, after which it decreases again(*4*), raising the possibility that an accelerated transdifferentiation of the small cells towards macrophages moves them out of the time window of highest responsiveness. We therefore tested the effect of a shorter pulse of C/EBPa (6h) and found that the small cells were still highly resistant to reprogramming, with an 18h pulse still producing a slight increase in the number of iPSC colonies (Fig. S9E). This result indicates that the two cell types exhibit intrinsic differences in their cell conversion preferences.

We next asked whether the Myc signature also predicts the reprogramming efficiency of other somatic cells, examining existing datasets of 20 hematopoietic and non-hematopoietic cell types(*2, 17–20*). Of note, our analysis revealed that Myc signature expression in the starting cell types is strongly predictive of the reprogramming efficiency (R=0.66, p=0.0016). In support of our findings in B cells, myeloid progenitors (MPs) and GMPs exhibited the highest levels of the Myc signature and the highest reprogramming efficiency (Fig.3J).

In summary, we have discovered by single cell RNA sequencing that two somatic cell types, only a single differentiation step apart, differ substantially and reciprocally in their propensity to transdifferentiate or reprogram. Whereas large pre-BII cells are highly susceptible for reprogramming into iPS cells and transdifferentiate slowly into macrophages, the small pre-BII cells, into which they normally differentiate, reprogram less efficiently but transdifferentiate more rapidly (Fig. 4).

**Fig. 4.**
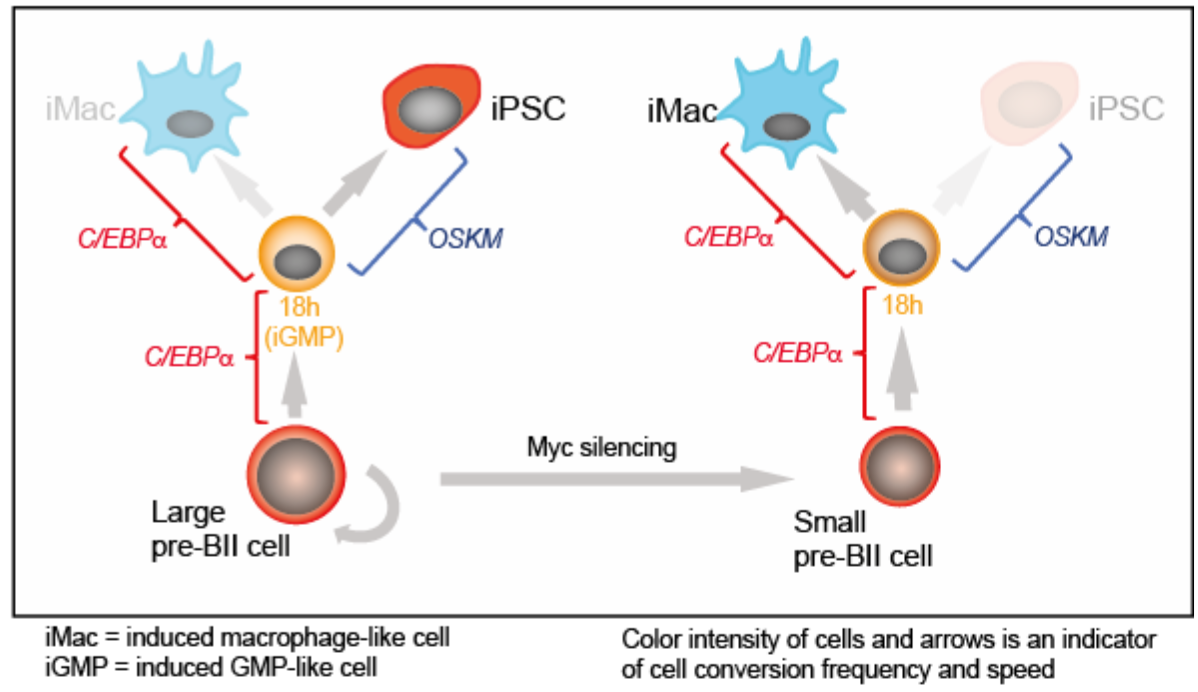
**Summary of the main findings.**

Our findings show that a cell’s propensity for either transdifferentiation or reprogramming can be dissociated, suggesting that these two types of plasticity might be fundamentally different. Our observations also suggest a link between Myc levels and cell identity. Specifically, B cells with high Myc levels are strongly biased for the acquisition of the pluripotent stem cell fate, while cells with low Myc levels transdifferentiate more rapidly. Accordingly, high levels of Myc are predictive of the iPSC reprogramming efficiency of diverse progenitor and mature cell types. The effect of Myc could be mediated by the factor’s ability to induce cell proliferation(*21*), global chromatin changes(*22, 23*), transcriptional amplification of genes essential for proliferation(*24*), changes in metabolism(*21*) and others. These features are also likely central for its role as a major driver of cancer(*21*) and its role in early embryonic development(*25*). However, high Myc expression in large pre-BII cells is not sufficient to enable their efficient iPSC reprogramming induced by OSKM, as we found that they must still be primed by C/EBPa(*4*). This might be related to C/EBPa’s ability to activate key transcription factors (such as Klf4 and Tet2), to recruit chromatin related factors (such as LSD1/Kdm1a, Hdac1, and Brd4)(*6*), and/or to induce changes in genome topology preceding pluripotent transcription factor expression(*13*). It is therefore tempting to speculate that Myc and a transiently expressed lineage instructive transcription factor such as C/EBPa are key ‘priming’ ingredients for the formation of pluripotent stem cells during cell reprogramming and normal early embryo development.

## ACKNOWLEDGEMENTS

We would like to thank Ido Amit and Diego Adhemar Jaitin for help with the MARS-Seq technique and the CRG/UPF FACS Unit for help with the cell sorting. Work in the lab of T.G. was supported by the European Research Council (ERC) Synergy Grant (4D-Genome) and by AGAUR (SGR-1136). Research in the lab of B.L. was supported by an ERC Consolidator grant (616434), the Spanish Ministry of Economy and Competitiveness (BFU2011-26206), the AXA Research Fund, the Bettencourt Schueller Foundation and AGAUR (SGR-831). We acknowledge support of the Spanish Ministry of Economy and Competitiveness, Centro de Excelencia Severo Ochoa 2013-2017 (SEV-2012-0208) and of the CERCA Programme / Generalitat de Catalunya.

## AUTHOR CONTRIBUTIONS

T.G., B.D.S., B.L. and M.F. conceived the study and wrote the manuscript. T.G. and B.L. supervised the work. B.D.S., C.B. and M. de A. performed reprogramming experiments and FACS analyses. M.F. performed all bioinformatics analyses and G.R.E. the pre-analyses of the data. M.M.L., A.G., H.H., I.G., and M.G. set up the single MARS-Seq technology.

## Supplementary materials

### Materials and methods

#### Mice and cell cultures

We used ‘reprogrammable mice’ containing a doxycycline-inducible OSKM cassette and the tetracycline transactivator(*5*). CD19^+^ pre-B cells were isolated from the bone marrow of these mice using monoclonal antibody to CD19 (clone 1D3, BD Pharmingen #553784) and MACS sorting (Miltenyi Biotech). Cell purity was confirmed to be >98% CD19+ by FACS using an LSRII machine (BD). After isolation, B cells were grown in RPMI medium supplemented with 10% FBS and 10ng/ml IL-7 (Peprotech), L-glutamine, nonessential amino acids, β-mercaptoethanol (Life Technologies) as well as penicillin/streptomycin. Mouse embryo fibroblasts (MEFs) were isolated from E13.5 mouse and expanded in DMEM supplemented with 10% FBS, L-glutamine and penicillin/streptomycin. Cultures were routinely tested for mycoplasma contamination. Animal experiments were approved by the Ethics Committee of the Barcelona Biomedical Research Park (PRBB) and performed according to Spanish and European legislation.

#### Transdifferentiation and reprogramming experiments

For transdifferentiation pre-B cells were infected with C/EBPαER-hCD4 retrovirus produced by the PlatE retroviral packaging cell line (Cell Biolabs, # RV-101). The cells were expanded for 48hrs on Mitomycin C-inactivated S17 feeders grown in RPMI medium supplemented with 10 ng/mL each of IL-7 (Peprotech) and hCD4^+^ were sorted (FACSaria, BD). For transdifferentiation C/EBPa was induced by treating the cells with 100nM β-Estradiol (E2) in medium supplemented with 10 ng/mL each of IL-7, IL-3 (Peprotech) and human colony-stimulating factor 1 (hCSF-1, kind gift of E. Richard Stanley). For reprogramming hCD4^+^ cells were plated at 500 cells/cm^2^ in gelatinized plates (12 wells) on irradiated MEF feeders in RPMI medium and pre-treated for 18h with E2 to induce C/EBPα. After E2 washout the cultures were switched to serum-free N2B27 medium supplemented with 10ng/ml IL-4, IL-7 and IL-15 (Peprotech) at 2ng/ml and treated with 2μg/ml of doxycycline to activate OSKM. From day 2 onwards the N2B27medium was supplemented with 20% KSR (Life Technologies), 3uM CHIR99021 and 1uM PD0325901 (2i medium). A step-by-step protocol describing the reprogramming procedure can be found at Nature Protocol Exchange (https://www.nature.com/protocolexchange/protocols/4567).

#### Myc expression by flow cytometry

CD19 positive B cells were washed and fixed in Fix&Perm fixative (Life Technologies) for 15 min, then washed and permeabilized in Fix&Perm saponin-based permeabilization buffer for 15 min. After permeabilization, cells were incubated in 1× PBS / 10% normal goat serum / 0.3M glycine to block non-specific protein-protein interactions followed by Myc antibody at 1/76 dilution for 30 min at room temperature. The secondary antibody used was Goat Anti-Rabbit IgG H&L (Alexa Fluor^®^ 647) (Life technologies) at 1/2000 dilution for 30 min. A rabbit IgG was used as the isotype control. Cells were analysed on a BD LSRII flow cytometer. The gating strategy is described in Fig. S10A.

#### Cell cycle analysis by EdU incorporation

For cell cycle analyses cells were treated for 2 hrs with EdU (Life Technologies). EdU staining was performed using the Click-IT EdU Cytometry assay kit (Life Technologies) at room temperature following the manufacturer’s instructions. Briefly, cells were washed in PBS and fixed in Click-iT fixative for 15 min. After washing they were permeabilized in 1 × Click-iT saponin-based permeabilization buffer for 15 min. The EdU reaction cocktail (PBS, CuSO_4_, Alexa Fluor 488 azide and buffer additive as per manufacturer’s protocol) was added to the cells for 30 min and then washed in 1% BSA/PBS. After staining, cells were analysed on a BD LSRII flow cytometer. The gating strategy is described in Fig. S10B.

#### FACS analyses of transdifferentiation

B cell to macrophage transdifferentiation was monitored by flow cytometry using antibodies against Mac-1 (clone 44, BD Pharmingen) and CD19 (1D3, BD Pharmingen) labeled with APC and PeCy-7, respectively. After staining, cells were analysed on a BD LSRII flow cytometer. The gating strategy is described in Fig. S10C.

#### RNA extraction

To remove the feeders, cells were trypsinized and pre-plated for 30min before RNA isolation with the miRNeasy mini kit (Qiagen). RNA was eluted from the columns using RNase-free water and quantified by Nanodrop. cDNA was produced with the High Capacity RNA-to-cDNA kit (Applied Biosystems).

#### qRT-PCR analyses

qRT-PCR reactions were set up in triplicate with the SYBR Green QPCR Master Mix (Applied Biosystems). Reactions were run on an AB7900HT PCR machine with 40 cycles of 30s at 95 °C, 30s at 60 °C and 30s at 72 °C.

#### Viral vector and infection

Production of the C/EBPαER-hCD4 retroviral vector and B cell infection were performed as before(*4, 6*).

#### Alkaline Phosphatase (AP) staining

AP staining was performed using the Alkaline Phosphatase Staining Kit (STEMGENT) following the manufacturer’s instructions.

##### Library preparation and sequencing

Single-cell libraries from polyA-tailed RNA were constructed applying massively parallel single-cell RNA sequencing (MARS-Seq) (*7*) as described in (*27*). Single cells were FACS isolated into 384-well plates with lysis buffer and reverse-transcription primers containing the single-cell barcodes and unique molecular identifiers (UMIs). Each library consisted of two 192 single-cell pools per time point (pool a and pool b). Multiplexed pools were sequenced in an Illumina HiSeq 2500 system. Primary data analysis was carried out with the standard Illumina pipeline following the manufacturer’s protocol.

##### Data pre-processing

Quality check of sequenced reads was performed with the FastQC quality control tool(*28*). Samples that reached the quality standards were then processed to deconvolute the reads to cell level by de-multiplexing according to the pool and the cell barcodes, wherein the first read contains the transcript sequence and the second read the cell barcode and the UMI.

Samples were mapped and gene expression was quantified with default parameters using the RNA pipeline of the GEMTools 1.7.0 suite(*29*) on the mouse genome assembly GRCm38 (*30*) and Gencode annotations M8(*31*). We took advantage of the UMI information to correct for amplification biases during the library preparation, collapsing read counts for reads mapping on a gene with the same UMI and considering unambiguously mapped reads only.

#### Data analysis

Cells with a library size < 1800 were excluded from further analysis. Genes detected in less than 50 cells or less than 15 cells per group were also excluded from further analysis, resulting in expression data for 17183 genes in 3152 cells. Size factor normalization was applied by dividing the expression of each gene in each cell by the total number of detected mRNA molecules and multiplying by the median number of molecules across cells. An inverse hyperbolic sine transformation (log (x+sqrt(x^2^+1)), where x is the mRNA count) was then applied and the data was subsequently centred.

#### Dimensionality reduction, batch correction and gene expression reconstruction

We performed principal component analysis (PCA) by computing partial singular value decomposition (SVD) on the data matrix extracting the first 100 largest singular values and corresponding vectors using the method implemented in R in the ‘irlba’ package(*32*). The distribution of the singular values flattens out after 35 components (Fig. S1B). Examining singular vectors highlights the presence of batch effects between the two pools at each time point starting from component 3 (Fig. S1C). We therefore applied a batch correction method based on finding mutual nearest neighbours between batches (*33*). We used the R implementation (function ‘mnn’ in the ‘scran’ package) with k=15 nearest neighbours, and computing the nearest neighbours on the first 2 PCA dimensions which only capture biological variation. This method corrects batch effects shared across all samples. However, partial SVD on batch corrected data shows that among the first 35 components that retain signals (Fig. S1D) batch effects between the two pools are still present (Fig. S1E). We therefore applied independent component analysis (ICA) to decompose expression into 35 mutually independent components and estimate the relative mixing matrix that, when multiplied by the independent components, results in the observed data. ICA separates well sample-specific batch effects from biological signal (Fig. S1F). We filtered out components when the interquartile ranges of the distributions of component scores of the two pools do not overlap at any time point (components 3, 9, 13, 15, 16, 17, 19, 20, 21, 24, 26, 27,32, 35). A component correlated with cell position in the plate (Component 33, Fig. S1G) was also filtered out. We then reconstructed gene expression by multiplying filtered gene loadings (Table S1) by the filtered samples scores (Table S2) including only the selected 20 components (Fig. S2). The resulting gene expression matrix was then normalized using quantile normalization.

##### Characterization of the components: Gene set enrichment analysis

We clustered genes according to the loadings on the components from ICA, assigning each gene to the component with highest or lowest loading. Each component therefore defines one cluster of negatively correlated genes and one of positively correlated genes. We then calculated the enrichment of each cluster for Gene Ontology categories(*34*), restricting the analysis to categories including more than 10 and less than 200 genes, and hallmark signatures from the Molecular Signature database(*35*) and tested its significance using Fisher’s test. P-values were corrected for multiple testing using Benjamini-Hochberg method(*36*).

##### Characterization of the components: comparison to the mouse cell atlas

We compared our data to a comprehensive atlas of murine cell types (*26*). We applied ICA to decompose expression of the atlas of cell types into 120 mutually independent components, and we correlated these to the components extracted from our single cell data (Fig. S2) to determine cell type specificity of single cell components.

##### Diffusion map

To visualize data in low dimensional space we used diffusion maps. Diffusion maps are a method for non-linear dimension reduction that learn the non-linear data manifold by computing the transition probability of each data point to its neighbours (diffusion distances). We used the R implementation by (*37*) available in package ‘dpt’ version 0.6.0. The transition matrix is calculated on the selected ICA components using a sigma = 0.12 for the Gaussian kernel.

##### Computation of similarity index of our single cell RNA-seq data with reference cell types

We compared our data to a comprehensive atlas of murine cell types from(*26*) that consists of uniformly re-analysed bulk and single cell RNA-seq data from 113 publications including 921 biological samples consisting of 272 distinct cell types.

We calculated a similarity index for each single cell transcriptome to each atlas cell type transcriptome as follows: we first calculated the genome wide correlation between each single cell and all cell types from the atlas. The correlation coefficient was then transformed using Fisher’s z transformation: 1/2 *log(1+r/1−r)). The vector of z-transformed correlations for each single cell was then scaled across reference cell types. In the same manner, we also compared our starting population single cell data to reference bulk expression data from different stages of B cell development from (*14*) and from the immunological genome project(*16*). Myc targets component increases in expression with time during reprogramming. This may fully account for the prediction of the extent of reprogramming in each cell. We therefore regress out the expression of Myc targets before the computation of similarity indices by removed the ‘Myc targets’ component from both the atlas and the single cell data before reconstructing both the atlas and single cell expression as explained above to derive a corrected similarity index. This shows that Myc targets are still predictive progression towards pluripotency at least at D4 (Fig. S8).

##### Correlation between reprogramming efficiency and the Myc component

Reprogramming efficiency data for different hematopoietic cell types as well as from mouse tail fibroblasts are from reference (*18*); for neural stem cells, pancreatic beta cells, keratinocytes and MEFs are from references (*20*) (*19*) (*17*) and (*2*), respectively. Cell reprogramming efficiencies were matched to the expression values of their Myc component, obtained from the mouse cell atlas (*26*) as described above (Fig. S2A, C)

When more than one cell type from the atlas corresponded to a single cell category used for reprogramming, their Myc component values were averaged (Table S5).

#### Data availability

Single cell gene expression data have been deposited in the National Center for Biotechnology Information Gene Expression Omnibus (GEO) under accession number GSE112004.

### Supplementary figures

**Fig. S1.**
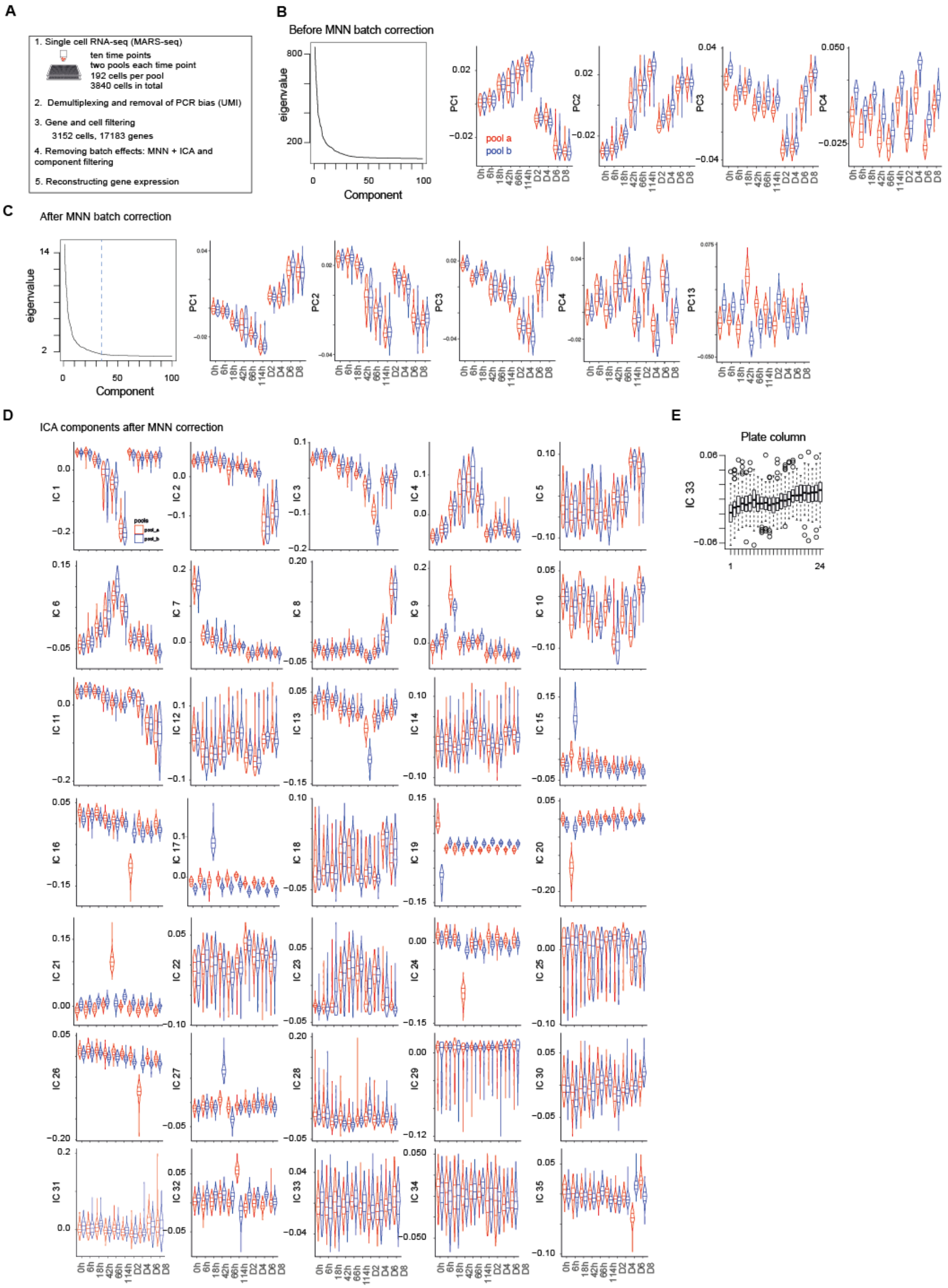
Data pre-processing, batch correction and independent component analysis. **A,** Overview of the data collection and pre-processing steps. **B,** Distribution of the top 100 eigenvalues and of single cell projections onto the first four principal components across pools and time points from the gene expression PCA before batch correction. **C,** Distribution of the top 100 eigenvalues and of single cell projections onto the first four principal components and component 13 across pools and time points from the PCA of gene expression after MNN batch correction. **D,** Distribution of single cells projections onto the 35 independent components across pools and time points. **E**, Distribution of the single cells projections onto independent component 33 across plate columns.

**Fig. S2.**
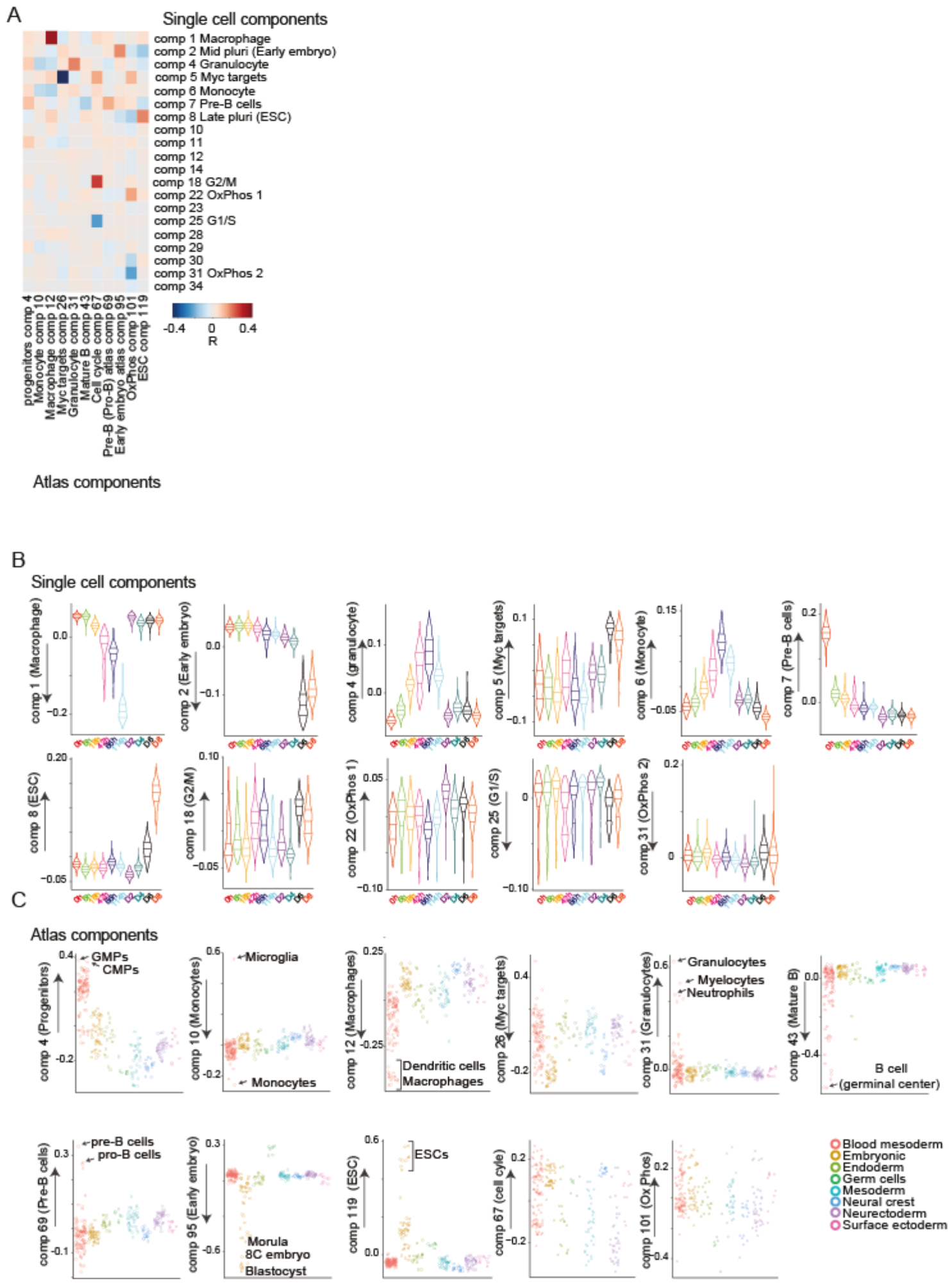
Characterization of independent components. **A,** Heatmap of the correlations between the gene loadings of selected single cell independent components and gene loadings of selected independent components from the reference mouse cell atlas(*26*). **B**, Distribution of the single cell projections onto the macrophage, mid pluripotency, granulocyte, monocyte, pre-B, late pluripotency, G2/M, oxidative phosphorylation, G1/S and a second oxidative phosphorylation specific components across time points. **C**, Cell type projections onto selected Atlas components.

**Fig. S3.**
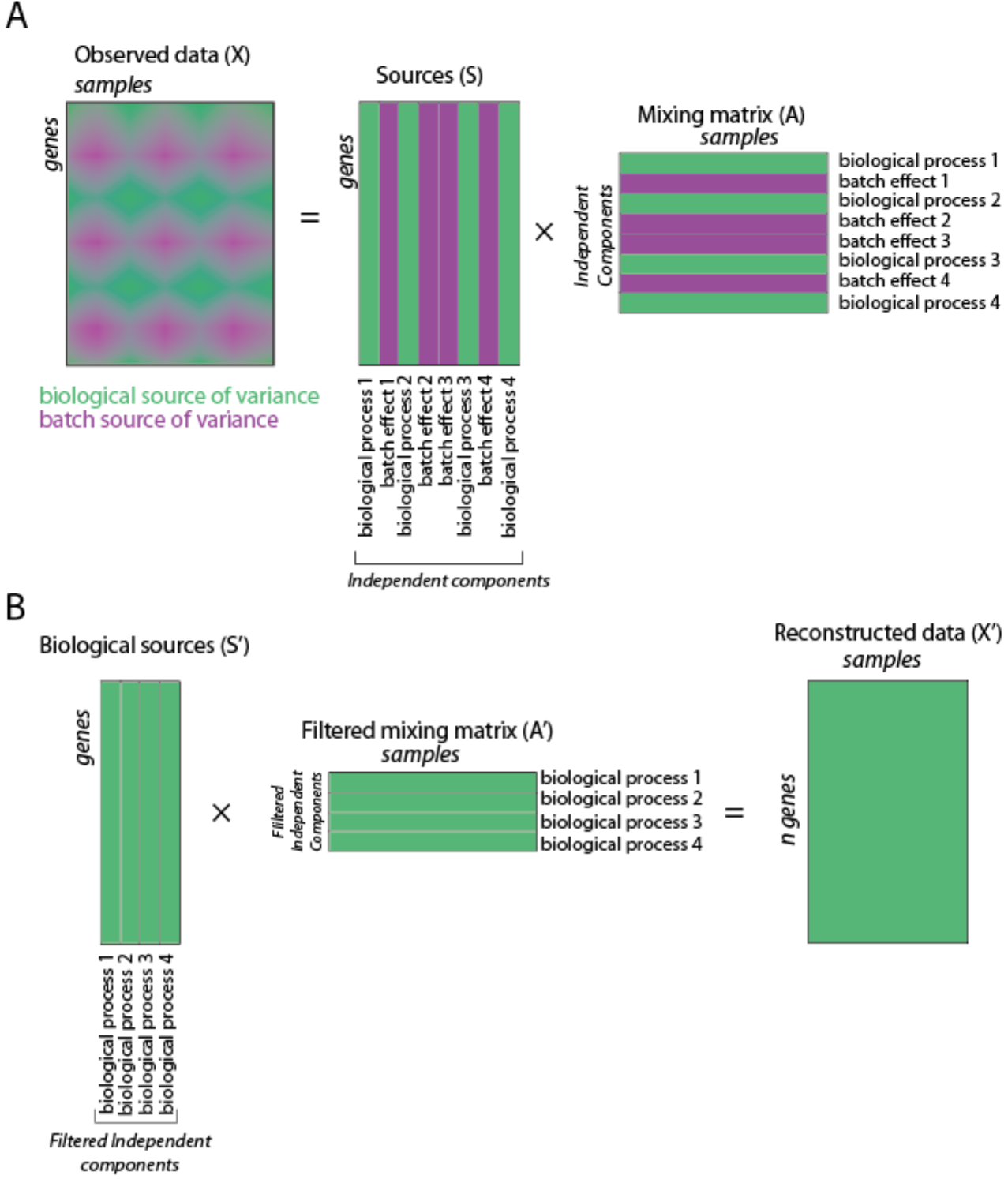
Reconstructing batch corrected gene expression. **A,** ICA decomposition the expression data matrix into a matrix of independent sources and mixing matrix. **B**, Reconstruction of gene expression after filtering out components capturing batch effects.

**Fig. S4.**
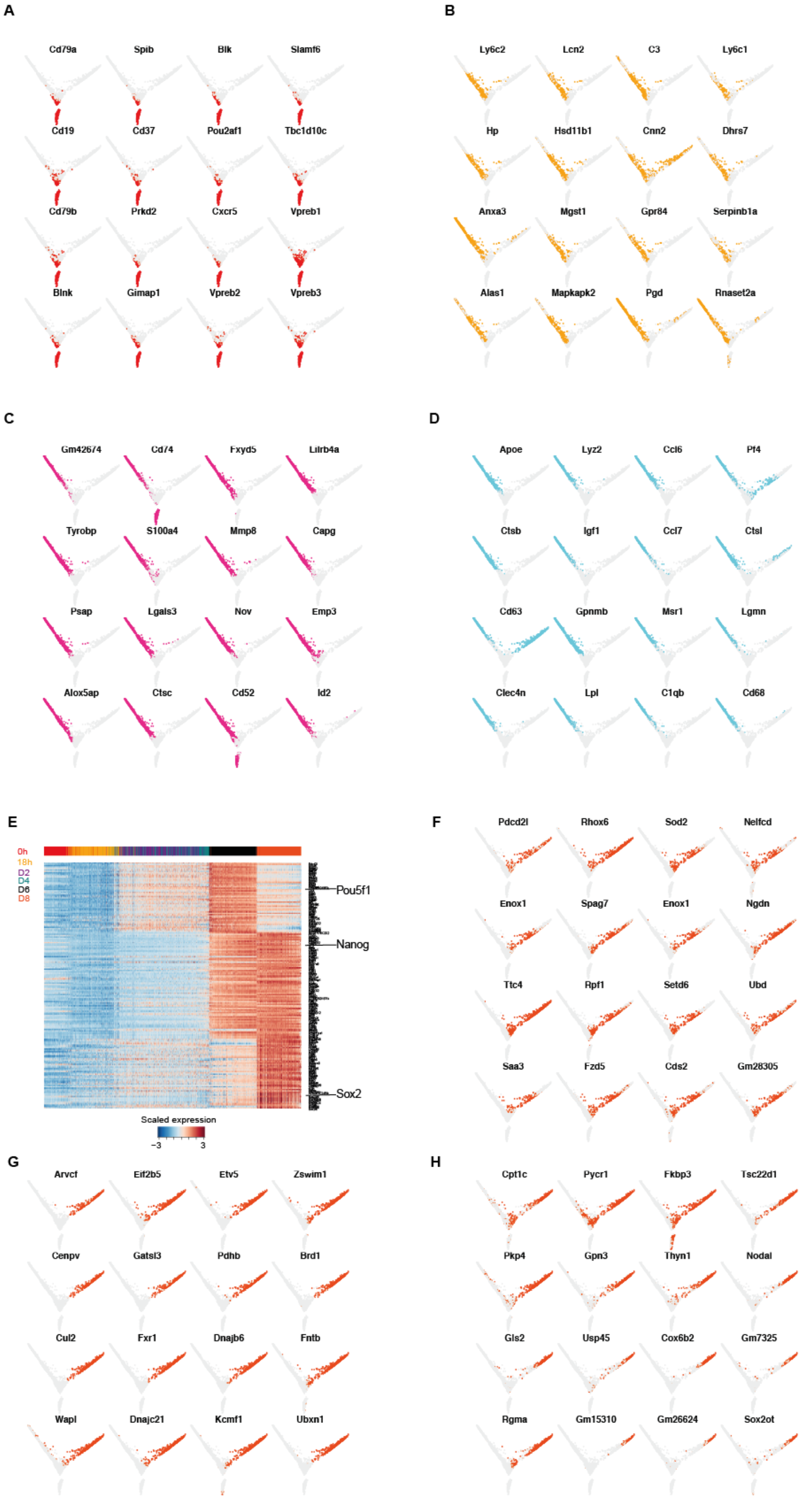
Single cell analysis of reprogramming and transdifferentiation. **A-D** Single cell projections onto the first two diffusion components, with cells expressing top 50% of selected markers for B cells in red (A), GMPs/granulocytes in light orange (**B**), monocytes in purple (**C**) and macrophages in light blue (**D**). **E**, Heatmap of genes up-regulated early (Pou5f1 cluster), mid (Nanog cluster) and late (Sox2 cluster) during reprogramming. **F-H**, Single cell projections onto the first two diffusion components, with cells expressing top 50% with of selected early (**F**), mid (**G**) and late (**H**) pluripotency markers in orange-red.

**Fig. S5.**
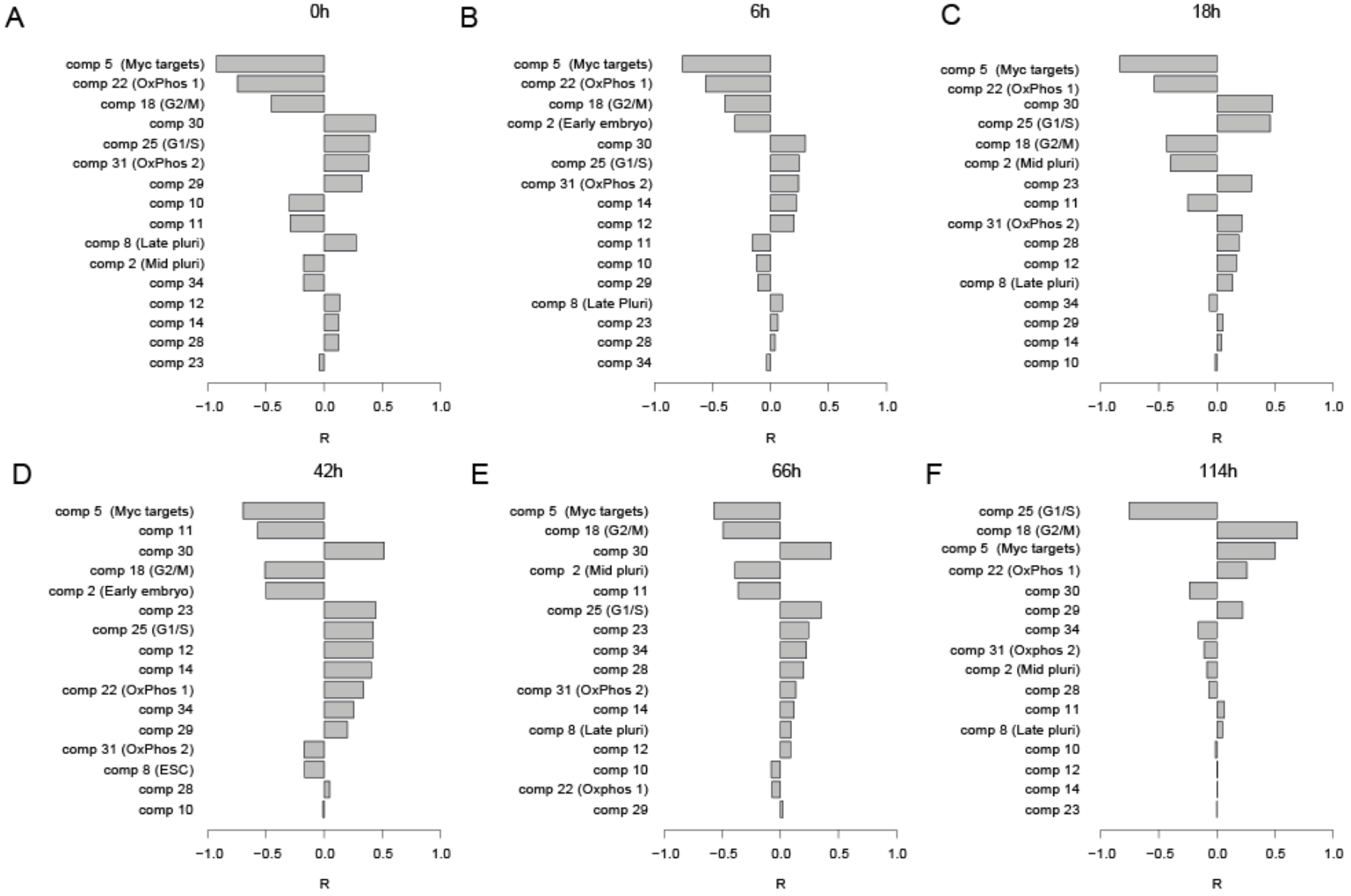
A-F. Predicting the speed of transdifferentiation. Correlation between each independent component and the expression similarity of single cells with reference bone marrow derived macrophages at 0h (**A**), 6h (**B**), 18h (**C**), 42h (**D**), 66h (**E**) and 114h (**F**) after C/EBPα induction.

**Fig. S6.**
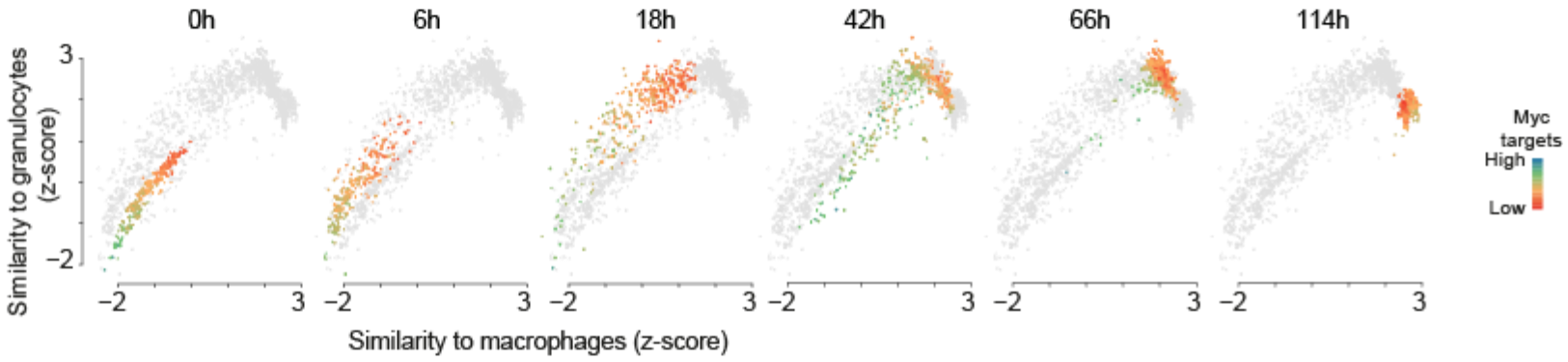
Acquisition of a transient granulocyte-like state during transdifferentiation. Single cell trajectories showing the relationship between granulocyte similarity and acquisition of the macrophage state during transdifferentiation. Colours indicate the levels of Myc targets.

**Fig. S7.**
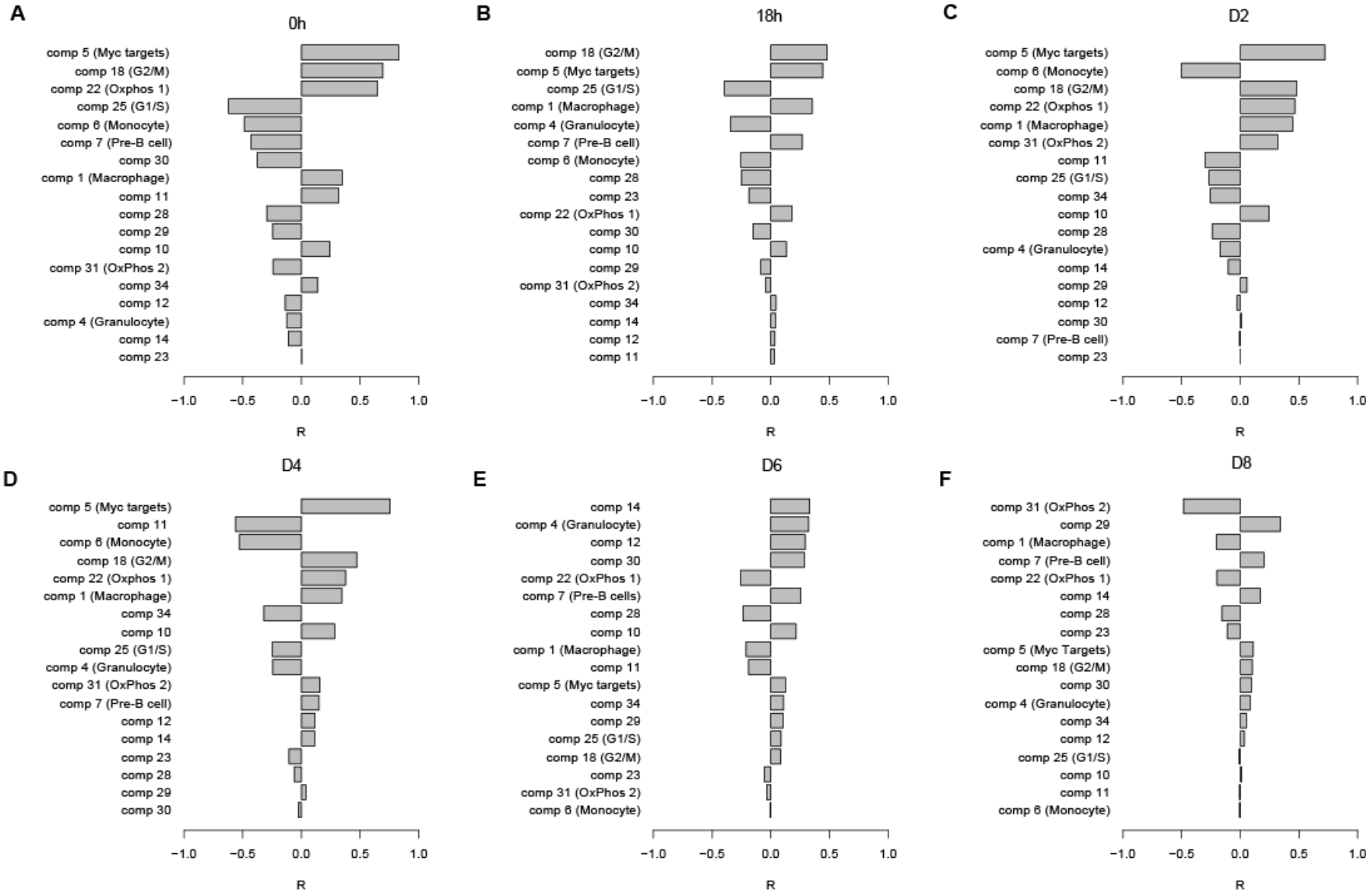
A-F. Predicting the speed of reprogramming. Correlation between each independent component and the expression similarity of single cells with acquisition of pluripotency at 0h (**A**) and 18 hours after C/EBPa induction (**B**), and at D2 (**C**), D4 (**D**), D6 (**E**) and D8 (**F**) after OSKM induction.

**Fig. S8.**
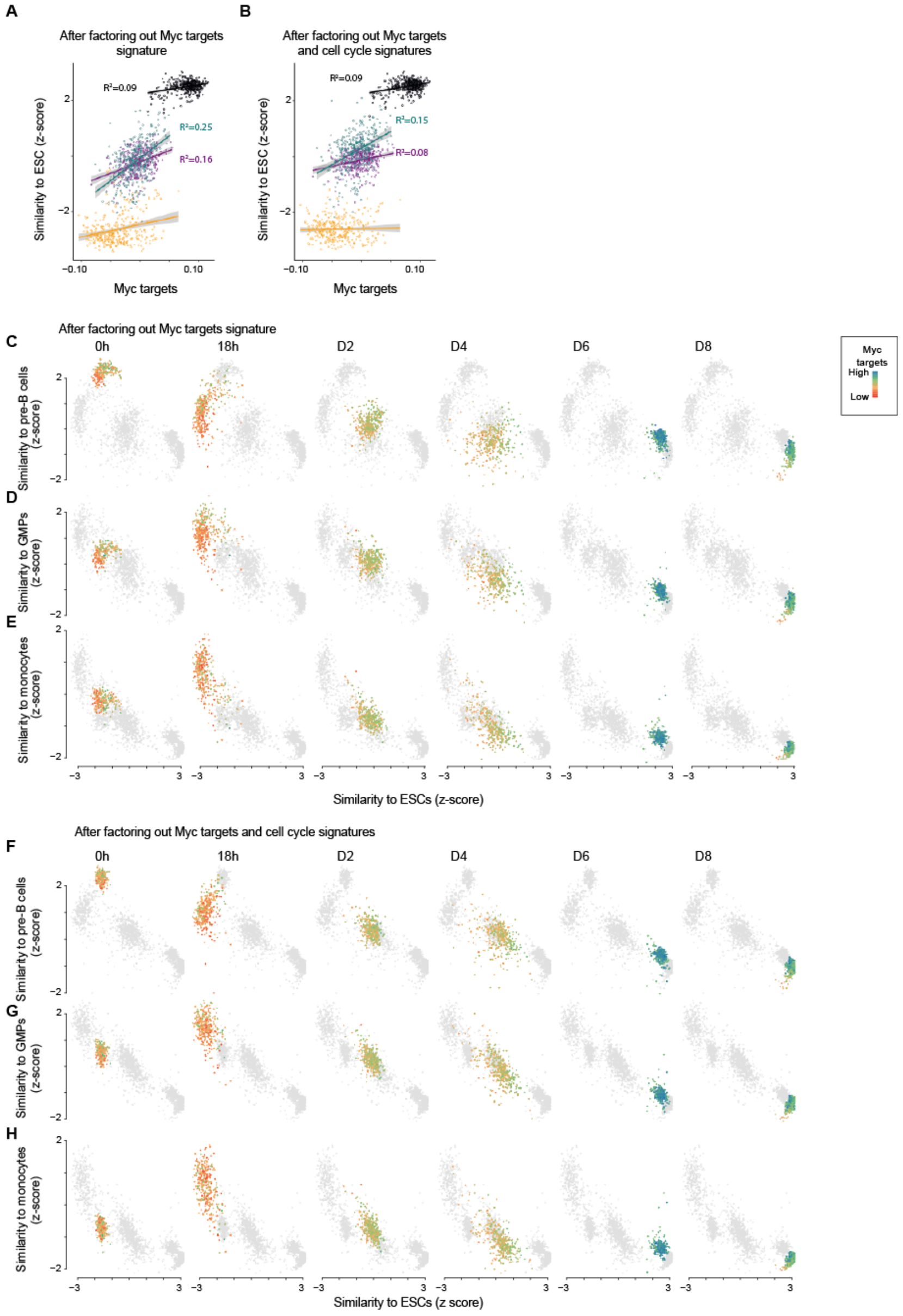
High expression of Myc targets predicts faster route towards reprogramming also when factoring out Myc targets and cell cycle components before the computation of the similarity score. **A,B** Correlation between Myc targets component and expression similarity of single cells to reference ESCs (acquisition of pluripotency) at each time point during reprogramming, calculated after factoring out Myc targets component (**A**) and both Myc and cell cycle components (**B**) from both single cell and Cell Atlas gene expression data (see Materials and Methods). **C**-**E** Loss of the B cell (**C**), GMP (**D**), and monocyte (**E**) state in relation to acquisition of pluripotency (calculated as in **A**) at each time point during reprogramming. **F-H,** Loss of the B cell (**F**), GMP (**G**), and monocyte (**H**) state in relation to acquisition of pluripotency (calculated as in **B**) at each time point during reprogramming. Colours indicate the levels of Myc targets.

**Fig. S9.**
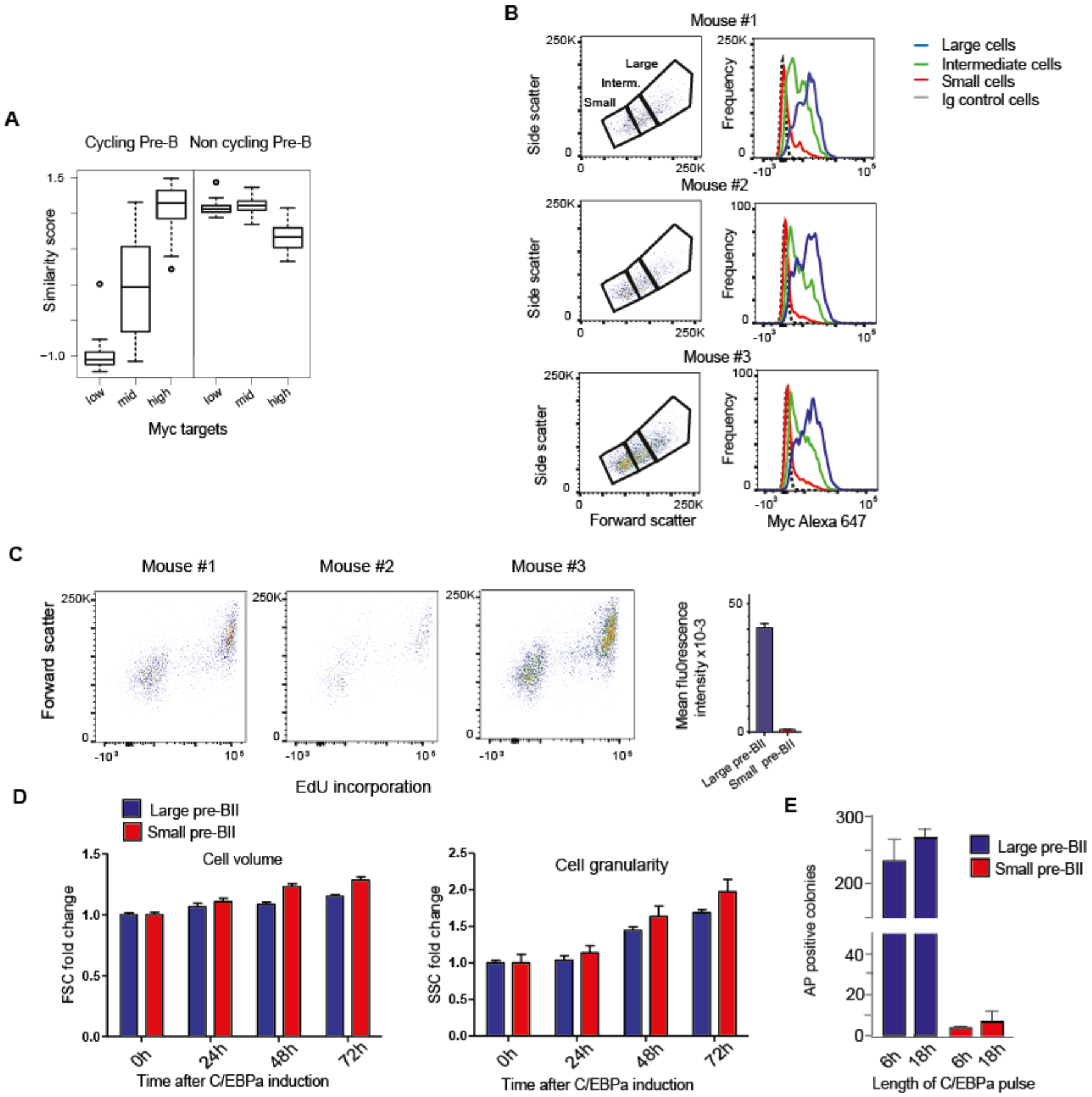
Experimental data relevant for Fig. 3. **A,** Similarity score of single cells binned by Myc targets expression (bottom 20%, mid and top 20%) with reference cycling and non-cycling pre-B cells(*16*). **B,** Top: FACS plots from pre-B cells obtained from 3 separate mice, showing the distribution of cells by volume (FSC) and granularity (SSC). Bottom: Myc expression profiles obtained for large, intermediate and small cells (gated in the profiles on the top) after intracellular immunostaining and FACS analysis. **C,**Cell proliferation analysis by FACS of uninduced pre-B cells by EdU incorporation for 2 hours. **D,**Monitoring cell volume and granularity during induced transdifferentiation of large and small pre-BII cells by SSC and FSC. **E**, Number of AP^+^ iPSC colonies at day 12 of reprogramming, obtained from large and small pre-BII cells pre-treated for either 6h or 18h of C/EBPa induction.

**Fig. S10.**
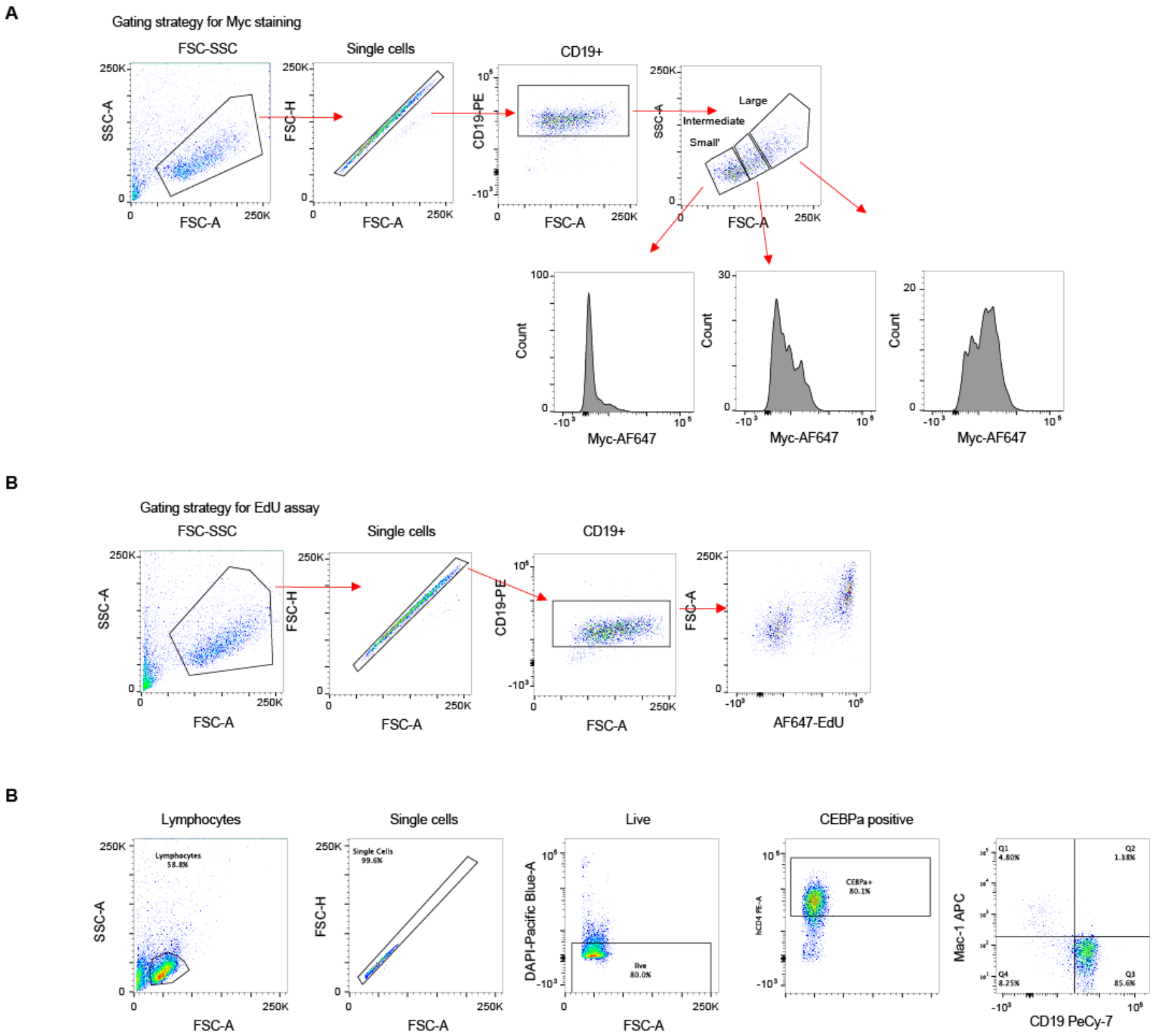
Gating strategies for FACS analyses. **A,** Gating strategy for Myc staining, corresponding to Fig. 3E and Fig. S9B. **B**, Gating strategy for EdU incorporation, corresponding to Fig. S9C. **C**, Gating strategy for transdifferentiation, corresponding to Fig. 3F.

### Supplementary tables

**Table S1.** Gene cluster membership and gene loadings on each independent component for each detected gene. The sign of cluster membership is positive if the gene has the highest absolute loading on the positive side of the component and negative if the highest absolute loading is on the negative side of the component.

**Table S2.** Total mRNA count, number of detected genes, and projection onto each independent component, for each single cell.

**Table S3.** Fisher’s test based gene set enrichment analysis on Gene Ontology categories (biological process) for each gene cluster. Includes odds ratios, p-values and FDR, number of genes associated to each category, number and names of genes included both in the cluster and in the category.

**Table S4.** Fisher’s test based gene set enrichment analysis on hallmark genesets for each gene cluster. Includes odds ratio, p-value and FDR, number of genes included in each category, number and names of genes both included both in the cluster and in the category.

**Table S5.** Reprogramming efficiencies for different cell types and their expression of Myc component from the mouse cell type atlas.

## References

1. H. Xie, M. Ye, R. Feng, T. Graf, Stepwise reprogramming of B cells into macrophages. Cell 117, 663–676 (2004).

2. K. Takahashi, S. Yamanaka, Induction of pluripotent stem cells from mouse embryonic and adult fibroblast cultures by defined factors. Cell 126, 663–676 (2006).

3. C. Nerlov, The C/EBP family of transcription factors: a paradigm for interaction between gene expression and proliferation control. Trends Cell Biol 17, 318–324 (2007).

4. B. Di Stefano et al., C/EBPalpha poises B cells for rapid reprogramming into induced pluripotent stem cells. Nature 506, 235–239 (2014).

5. B. W. Carey, S. Markoulaki, C. Beard, J. Hanna, R. Jaenisch, Single-gene transgenic mouse strains for reprogramming adult somatic cells. Nat Methods 7, 56–59 (2010).

6. B. Di Stefano et al., C/EBPalpha creates elite cells for iPSC reprogramming by upregulating Klf4 and increasing the levels of Lsd1 and Brd4. Nat Cell Biol 18, 371–381 (2016).

7. D. A. Jaitin et al., Massively parallel single-cell RNA-seq for marker-free decomposition of tissues into cell types. Science 343, 776–779 (2014).

8. L. Haghverdi, F. Buettner, F. J. Theis, Diffusion maps for high-dimensional single-cell analysis of differentiation data. Bioinformatics 31, 2989–2998 (2015).

9. B. Treutlein et al., Dissecting direct reprogramming from fibroblast to neuron using single-cell RNA-seq. Nature 534, 391–395 (2016).

10. D. Cacchiarelli et al., Aligning single-cell developmental and reprogramming trajectories identifies molecular determinants of reprogramming outcome. bioRxiv, (2017).

11. L. Guo et al., Resolution of Reprogramming Transition States by Single Cell RNA-Sequencing. bioRxiv, (2017).

12. G. Schiebinger et al., Reconstruction of developmental landscapes by optimal-transport analysis of single-cell gene expression sheds light on cellular reprogramming. bioRxiv, (2017).

13. R. Stadhouders et al., Transcription factors orchestrate dynamic interplay between genome topology and gene regulation during cell reprogramming. Nat Genet 50, 238–249 (2018).

14. R. Hoffmann, T. Seidl, M. Neeb, A. Rolink, F. Melchers, Changes in gene expression profiles in developing B cells of murine bone marrow. Genome Res 12, 98–111 (2002).

15. R. Nahar et al., Pre-B cell receptor-mediated activation of BCL6 induces pre-B cell quiescence through transcriptional repression of MYC. Blood 118, 4174–4178 (2011).

16. M. W. Painter et al., Transcriptomes of the B and T lineages compared by multiplatform microarray profiling. The Journal of Immunology 186, 3047–3057 (2011).

17. T. Aasen et al., Efficient and rapid generation of induced pluripotent stem cells from human keratinocytes. Nat Biotechnol 26, 1276–1284 (2008).

18. S. Eminli et al., Differentiation stage determines potential of hematopoietic cells for reprogramming into induced pluripotent stem cells. Nat Genet 41, 968–976 (2009).

19. M. Stadtfeld, K. Brennand, K. Hochedlinger, Reprogramming of pancreatic beta cells into induced pluripotent stem cells. Curr Biol 18, 890–894 (2008).

20. J. B. Kim et al., Pluripotent stem cells induced from adult neural stem cells by reprogramming with two factors. Nature 454, 646–650 (2008).

21. C. V. Dang, MYC on the path to cancer. Cell 149, 22–35 (2012).

22. P. S. Knoepfler et al., Myc influences global chromatin structure. EMBO J 25, 2723–2734 (2006).

23. K. R. Kieffer-Kwon et al., Myc Regulates Chromatin Decompaction and Nuclear Architecture during B Cell Activation. Mol Cell 67, 566–578 e510 (2017).

24. C. Y. Lin et al., Transcriptional amplification in tumor cells with elevated c-Myc. Cell 151, 56–67 (2012).

25. R. Scognamiglio et al., Myc Depletion Induces a Pluripotent Dormant State Mimicking Diapause. Cell 164, 668–680 (2016).

26. A. P. Hutchins et al., Models of global gene expression define major domains of cell type and tissue identity. Nucleic Acids Res 45, 2354–2367 (2017).

27. A. Guillaumet-Adkins et al., Single-cell transcriptome conservation in cryopreserved cells and tissues. Genome Biol 18, 45 (2017).

28. S. Andrews. (http://www.bioinformatics.babraham.ac.uk/projects/fastqc, 2010).

29. S. Marco-Sola, M. Sammeth, R. Guigo, P. Ribeca, The GEM mapper: fast, accurate and versatile alignment by filtration. Nat Methods 9, 1185–1188 (2012).

30. F. Cunningham et al., Ensembl 2015. Nucleic Acids Res 43, D662–669 (2015).

31. J. M. Mudge, J. Harrow, Creating reference gene annotation for the mouse C57BL6/J genome assembly. Mamm Genome 26, 366–378 (2015).

32. J. Baglama, L. Reichel, Augmented implicitly restarted Lanczos bidiagonalization methods. SIAM Journal on Scientific Computing 27, 19–42 (2005).

33. L. Haghverdi, A. T. L. Lun, M. D. Morgan, J. C. Marioni, Correcting batch effects in single-cell RNA sequencing data by matching mutual nearest neighbours. bioRxiv, (2017).

34. M. Ashburner et al., Gene ontology: tool for the unification of biology. The Gene Ontology Consortium. Nat Genet 25, 25–29 (2000).

35. A. Liberzon et al., The Molecular Signatures Database (MSigDB) hallmark gene set collection. Cell Syst 1, 417–425 (2015).

36. Y. Benjamini, Y. Hochberg, Controlling the false discovery rate: a practical and powerful approach to multiple testing. Journal of the royal statistical society. Series B (Methodological), 289–300 (1995).

37. L. Haghverdi, M. Buttner, F. A. Wolf, F. Buettner, F. J. Theis, Diffusion pseudotime robustly reconstructs lineage branching. Nat Methods 13, 845–848 (2016).

